# Comprehensive *SMN1* and *SMN2* profiling for spinal muscular atrophy analysis using long-read PacBio HiFi sequencing

**DOI:** 10.1101/2022.10.19.512930

**Authors:** Xiao Chen, John Harting, Emily Farrow, Isabelle Thiffault, Dalia Kasperaviciute, Genomics England Research Consortium, Alexander Hoischen, Christian Gilissen, Tomi Pastinen, Michael A Eberle

**Author notes:** Correspondence: Michael Eberle.

## Abstract

Spinal muscular atrophy, a leading cause of early infant death, is caused by biallelic mutations of the *SMN1* gene. Sequence analysis of *SMN1* is challenging due to high sequence similarity with its paralog *SMN2*. Both genes have variable copy numbers across populations. Furthermore, without pedigree information, it is impossible to identify silent carriers (2+0) with two copies of *SMN1* on one chromosome and zero copies on the other. We developed Paraphase, an informatics method that identifies full-length *SMN1* and *SMN2* haplotypes, determines the gene copy numbers and calls phased variants using long-read PacBio HiFi data. The *SMN1* and *SMN2* copy number calls by Paraphase are highly concordant with orthogonal methods (99.2% for *SMN1* and 100% for *SMN2*). We applied Paraphase to 438 samples across five ethnic populations to conduct a population-wide haplotype analysis of these highly homologous genes. We identified major *SMN1* and *SMN2* haplogroups and characterized their co-segregation through pedigree-based analyses. We identified two *SMN1* haplotypes that form a common two-copy *SMN1* allele in African populations. Testing positive for these two haplotypes in an individual with two copies of *SMN1* gives a silent carrier risk of 88.5%, which is significantly higher than the currently used marker (1.7-3.0%). Extending beyond simple copy number testing, Paraphase can detect pathogenic variants and enable potential haplotype-based screening of silent carriers through statistical phasing of haplotypes into alleles. Future analysis of larger population data will allow identification of more diverse haplotypes and genetic markers for silent carriers.

## Introduction

Spinal muscular atrophy (SMA) is a neuromuscular disease caused in most cases by biallelic mutations of the *SMN1* gene^1–3^. SMA is a leading cause of early infant death with an incidence of 1 in 6000-10,000 live births and a carrier frequency of 1 in 40-80 across ethnic groups^4–8^. SMA can be classified into four clinical types (type I-IV) that differ in age of onset and disease severity^1^.

*SMN1* and its paralog *SMN2* reside in a highly complex genomic region on chromosomal band 5q13 that is frequently subject to unequal crossing over and gene conversion, resulting in variable copy numbers (CNs) of *SMN1* and *SMN2*^7,9^. *SMN1* and *SMN2* are near-identical in sequence with just one functionally different base, NM_000344.3:c.840C>T. In *SMN2*, c.840T disrupts a splicing enhancer and leads to skipping of Exon 7^10^ and, as a result, most *SMN2* transcripts are unstable and almost nonfunctional. Since *SMN2* can produce a small amount of functional protein, the CN of *SMN2* is a modifier of the SMA disease severity^11^. The majority (∼96%) of 5q-linked SMA cases are caused by biallelic absence of *SMN1* c.840C through either large deletions or gene conversion to c.840T, while a smaller percentage (∼4%) are caused by other small pathogenic variants in *SMN1* in trans with c.840C loss^8,12–14^.

Because of the high carrier frequency and severity of SMA, the American College of Medical Genetics and Genomics recommends population-wide SMA screening^15^. Conventional SMA screening tests use PCR-based methods, such as multiplex ligation-dependent probe amplification (MLPA)^16,17^ and quantitative PCR (qPCR)^18^, to determine the *SMN1* dosage (copy number) in Exon 7, mostly targeting c.840C>T. To date, a few next-generation sequencing (NGS)-based *SMN1* callers have been reported^19–22^. These callers rely on short reads to identify copy number variations and distinguish *SMN1* and *SMN2* based on a limited number of differentiating bases centered around c.840C>T. However, dosage testing fails to identify carriers that carry pathogenic variants other than c.840C>T, which represent ∼1-2% of all carriers^5^. In addition, detecting *SMN2* variants in SMA patients is also important for understanding the disease modifying effect^23^. The *SMN1* or *SMN2* gene is ∼28kb long and detailed sequence analysis of the complete genes is labor-intensive for traditional Sanger sequencing and impossible for conventional short-read NGS methods due to the high sequence similarity between the two genes.

Furthermore, current tests (i.e. dosage testing) are unable to accurately phase alleles. This is important to distinguish between individuals carrying the normal *SMN1* genes on both alleles (1+1) and silent carriers (2+0) with two copies of *SMN1* on one chromosome and zero copies on the other. Silent carriers account for approximately 3-9% of carriers in non-African populations and 27% of carriers in African populations^5,6,21^. Throughout this paper, we use the term “singleton *SMN1* allele” to refer to chromosomes with a single copy of *SMN1*, and “two-copy *SMN1* alleles” to refer to alleles with two copies of *SMN1* occurring on the same chromosome. Previous studies have identified the g.27134T>G SNP (NM_000344.3:c.*3+80T>G) as a marker of the two-copy *SMN1* allele^24^ and this SNP is now commonly tested to modify the residual carrier risk - i.e. the probability that an individual with two copies of *SMN1* is a carrier. However, this SNP is rare and has low sensitivity in non-African populations. In Africans it is common but it is also present on almost 20% of singleton *SMN1* alleles^21^, so it does not have a high positive predictive value (PPV). When an African individual with two copies of *SMN1* tests positive for g.27134T>G, the residual risk of being a carrier, which is largely the silent carrier risk, is estimated to be just 1.7%-3.0%^20,21,24^. More population studies are needed to identify better markers to detect two-copy *SMN1* alleles, but again, short-read based methods suffer from the difficulty to differentiate *SMN1* from *SMN2* due to the high sequence similarity and thus are not ideal methods for identifying these markers.

To better facilitate SMA screening, there is an urgent need for a method that performs comprehensive full-gene *SMN1* and *SMN2* profiling. This method should ideally be able to 1) identify the CN of intact *SMN1* and *SMN2* based on c.840, 2) identify pathogenic variants in *SMN1* other than loss of c.840C, and 3) identify silent carriers. Accurate long-read sequencing is ideal for resolving regions with high sequence homology and the utility of long-read PacBio HiFi sequencing in *SMN1* was previously demonstrated in an amplicon-based study for a Chinese population^25^, though informatics methods are still lacking for shotgun HiFi sequencing, where high sequence homology results in ambiguous alignments. Here we describe a method, Paraphase, that accurately detects the CN, as well as variants throughout the *SMN1* and *SMN2* genes using PacBio HiFi sequencing. We applied Paraphase to population samples from five ethnicities and performed a population-wide haplotype analysis of these genes. We identified major haplogroups for *SMN1* and *SMN2* and quantified their co-segregation patterns. Furthermore, we identified specific haplotypes forming two-copy *SMN1* alleles which could greatly improve the accuracy of silent carrier detection.

## Materials and Methods

### Paraphase: HiFi-based *SMN1* and *SMN2* caller

Paraphase extracts HiFi reads aligned to either *SMN1* or *SMN2* and realigns them to the *SMN1* region. It then identifies variant positions throughout the 44kb long region of interest (chr5:70917100-70961220, GRCh38), which includes the *SMN1* gene body plus upstream/downstream regions. Paraphase then assembles haplotypes by linking the phases of each variant site (Figure 1). Haplotypes are assigned to *SMN1* or *SMN2* based on the sequence at the c.840 site, i.e. C is *SMN1* and T is *SMN2*. In addition, Paraphase identifies the common truncated form of *SMN2, SMN2Δ7–8* that has a 6.3kb deletion of Exons 7-8. Generally, the number of unique *SMN1* and *SMN2* haplotypes reflects *SMN1* and *SMN2* CNs. For samples with only one *SMN1* or *SMN2* haplotype identified, to rule out possible rare cases where two identical haplotypes exist, we calculate if the depth at the c.840C (T) site is consistent with one or two copies of *SMN1* (*SMN2*). A no-call is reported when the read depth could not reliably distinguish CN1 vs. CN2. CN calls are also adjusted when the number of supporting reads of one haplotype suggests twice the CN of the other haplotypes. With the complete haplotypes resolved, Paraphase makes phased variant calls throughout the genes by calling DeepVariant^26^. Paraphase also assigns haplotypes to haplogroups (see “Assigning haplotypes to haplogroups” section below) to enable further haplotype-based analysis for identifying genetic markers. Paraphase works on both whole-genome sequencing (WGS) or hybrid capture-based enrichment data.

**Figure 1.**
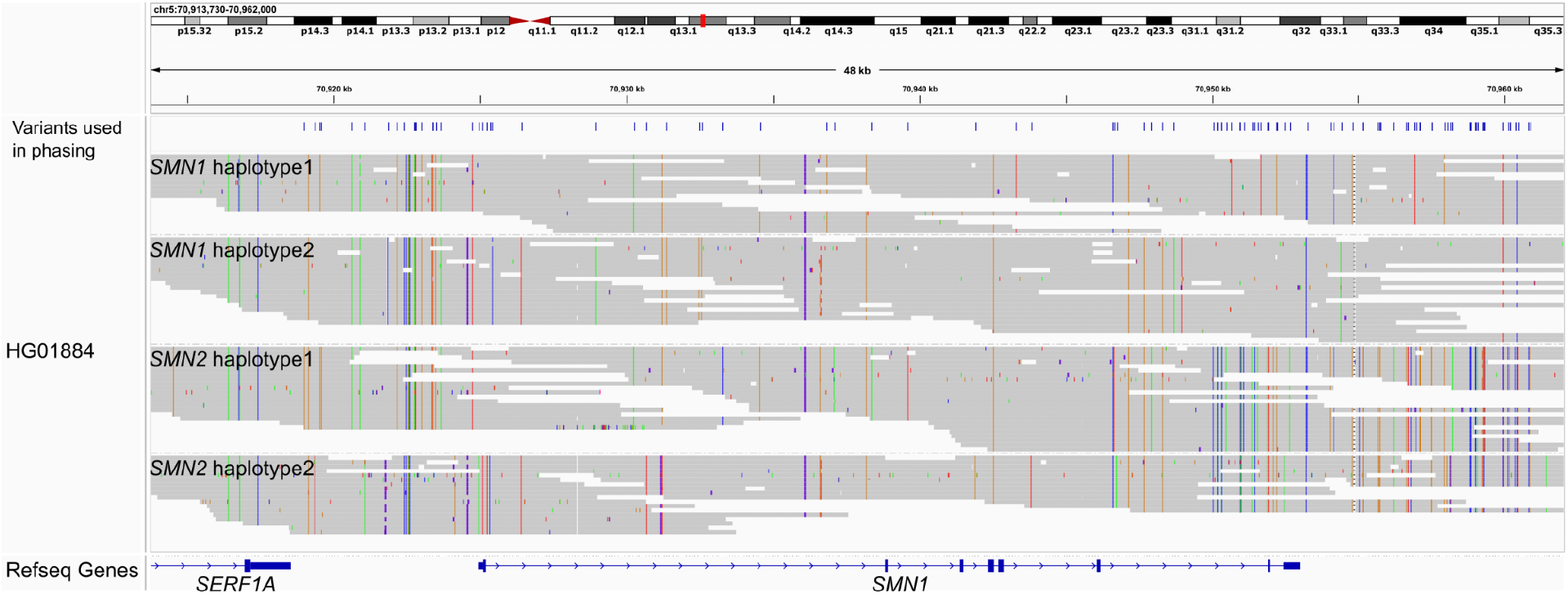
Visualization of assembled *SMN1* and *SMN2* haplotypes, taking HG01884 as an example. Paraphase produces haplotagged bamlets to facilitate examination of haplotypes with all relevant reads realigned to *SMN1*. Variant positions used in phasing are shown in the top panel and reads are grouped by their assigned haplotypes (IGV option: group by HP tag)

### Validation of CN calls

To verify the accuracy of our CN calls, we included 107 Coriell samples, 7 from Genome in a Bottle (GIAB)^27^ and 100 from the Human Pangenome Reference Center (HPRC)^28^. For these samples, *SMN1* and *SMN2* CNs were previously called by a short-read WGS based method^21^ which has been shown to have 99.7% concordance against MLPA and digital PCR. Three of the 107 samples had MLPA calls that agree with short-read based calls^29^. We also included 9 carrier (1+0) samples from Genomic Answers for Kids (GA4K) at Children’s Mercy Kansas City with MLPA results (SALSA MLPA P060 SMA Carrier probemix, MRC-Holland). Finally, we included an SMA trio from the 100,000 Genomes Project, where the *SMN1* CN of both parents are one and the proband has zero copy of *SMN1* (the *SMN2* CNs for these three samples are unknown). In total, we had 119 samples with *SMN1* CN information and 116 samples with *SMN2* CN information. Detailed validation sample information is summarized in Table S1.

### Population samples

We included 341 pedigrees (26 duos, 308 trios and 7 quartets) from five ethnic populations to study co-segregation of *SMN1* and *SMN2* alleles (Table S2 and Table S3). We collected these data from GIAB^27^, the Chinese Quartet project^30^, HPRC^28^, 1000 Genomes Project^31^, the 100,000 Genomes Project, Radboud University Medical Center and GA4K. Among these pedigrees, 198 are of European (EUR) origin, 37 African (AFR), 35 admixed American (AMR), 26 South Asian (SAS) and 18 East Asian (EAS), 18 of mixed ancestry and 9 of unknown ethnicity. In addition, we included 67 samples without pedigree information from GA4K for other frequency calculations (Table S3).

### Assigning haplotypes to haplogroups

Multiple sequence alignment and a neighbor-joining tree for the *SMN1* and *SMN2* haplotypes identified across populations were produced by Mafft server^32^ (version 7) with default parameters (https://mafft.cbrc.jp/alignment/server/). Haplogroups were identified by manually examining the tree for monophyletic groups. In Paraphase, a newly assembled haplotype in a given sample is assigned a haplogroup by comparing the sequence similarity with representative sequences from each haplogroup and selecting the most similar haplogroup. A small number of haplotypes from each haplogroup were used to produce trees in Figure 2 and Figure S1, visualized with FigTree v1.4.4 (https://github.com/rambaut/figtree/). Sequences of the same set of haplotypes were visualized in IGV in Figure 3.

**Figure 2.**
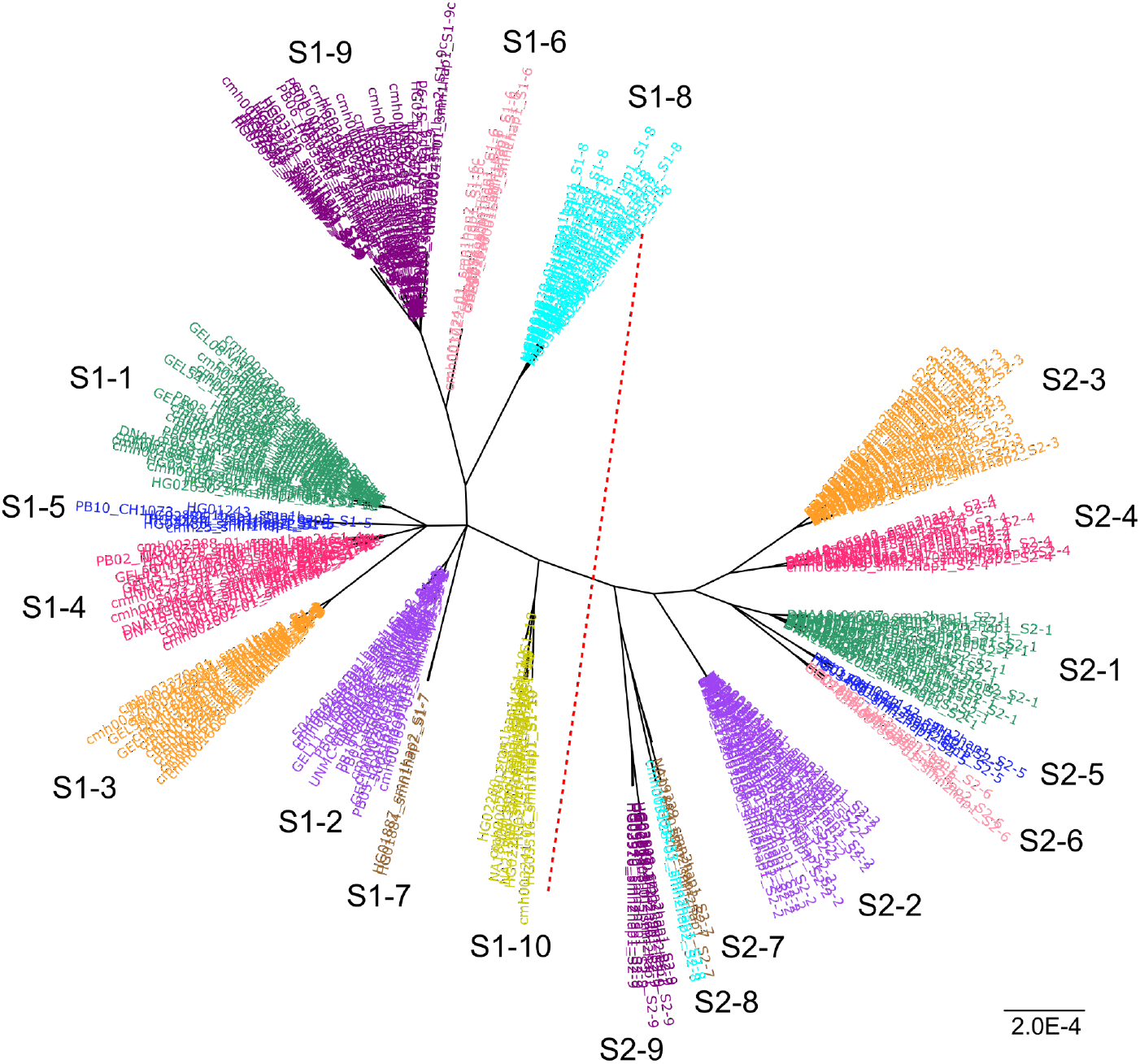
Population-wide haplotype analysis identified major *SMN1* and *SMN2* haplogroups. Representative haplotype sequences of the gene region from each *SMN1* and *SMN2* haplogroup were used to create an unrooted tree. The red dotted line in the middle separates *SMN1* (left) and *SMN2* (right). Figure S1 shows a tree of the same haplotypes created using the gene plus upstream/downstream regions, and a tree of the same haplotypes created using sequences of Exons 1-6.

**Figure 3.**
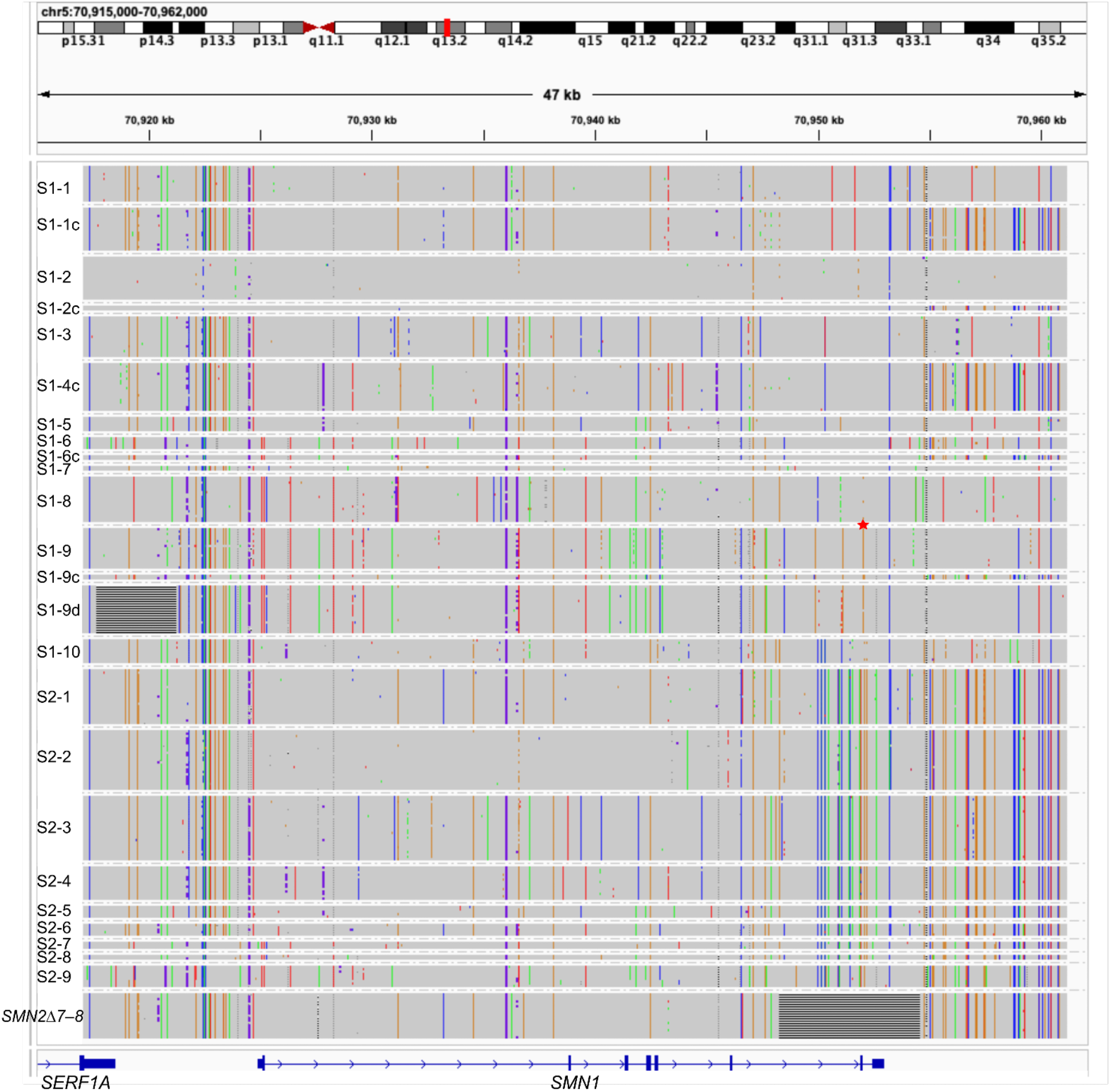
Representative haplotype sequences from each *SMN1* and *SMN2* haplogroup as well as *SMN2Δ7–8*. IGV snapshot showing the haplotype sequences used in the phylogenetic analysis in Figure 2. Sequences of the gene region plus upstream and downstream regions were included. *SMN2Δ7–8* has the 6.3kb deletion of Exons 7-8. S1-9d has a 3.6kb deletion upstream of *SMN1*. The SNP g.27134T>G, commonly used in silent carrier screening, is marked with a red star symbol between S1-8 and S1-9.

### Pedigree-based phasing of haplotypes into alleles

For this study, we use the term “haplotype” to refer to a set of phased variants (SNPs or indels) in one copy of a gene (*SMN1* or *SMN2*). Conversely, we use the term “allele” to refer to one or several haplotypes that are inherited on the same chromosome, e.g. co-segregation of two *SMN1* haplotypes or one *SMN1* and one *SMN2* haplotype. Phasing of haplotypes into alleles was done by comparing the haplotypes/haplogroups in parents and probands. Haplotypes were directly assigned haplogroups by Paraphase in samples with >20X HiFi WGS coverage. For parents with either Illumina short read data or low coverage HiFi data (Table S2), i.e. where phasing is not possible or accurate, representative variants for each haplogroup were queried in the parent data to identify the haplogroups in the parent.

Haplogroups carried by the parents and the proband were compared to identify which haplotype(s) is inherited on each allele. In ambiguous cases, i.e. both parents have haplotypes of the same haplogroup, manual examination of data in IGV was conducted to find unique SNPs that distinguish these haplotypes and phase them into alleles.

## Results

### Validation of Paraphase copy number calling

The *SMN1* and *SMN2* CN calls made by Paraphase are highly concordant with orthogonal methods, which include short-read WGS based CN calls, MLPA calls and SMA trio-based inference (see Methods). The CN call concordance is 99.2% for *SMN1* and 100% for *SMN2* (Table 1). We correctly called all SMA cases and carriers, and did not make any false positive case or carrier calls. The *SMN2Δ7–8* calls are also concordant with orthogonal methods.

**Table 1.** Validation against samples with known *SMN1*/*SMN2* copy numbers (CNs).

|  | CN by orthogonal methods | Total | Concordant | Discordant | No-call | Agreement (excluding no-calls) |
| --- | --- | --- | --- | --- | --- | --- |
| *SMN1* | 0 | 1 | 1 | 0 | 0 | 100% |
|  | 1 | 12 | 12 | 0 | 0 | 100% |
|  | 2 | 79 | 79 | 0 | 0 | 100% |
|  | >2 | 27 | 26 | 1<sup>a</sup> | 0 | 96.3% |
|  | Total | 119 | 118 | 1 | 0 | 99.2% |
| *SMN2* | 0 | 8 | 8 | 0 | 0 | 100% |
|  | 1 | 43 | 42 | 0 | 1<sup>b</sup> | 100% |
|  | 2 | 63 | 63 | 0 | 0 | 100% |
|  | >2 | 2 | 2 | 0 | 0 | 100% |
|  | Total | 116 | 115 | 0 | 1 | 100% |
| *SMN2*Δ7–8 | 0 | 104 | 104 | 0 | 0 | 100% |
|  | 1 | 3 | 3 | 0 | 0 | 100% |
|  | Total | 107 | 107 | 0 | 0 | 100% |

- The discordant call was a CN3 miscalled as CN2, due to two of the three haplotypes being identical in sequence. - The no-call was due to an ambiguous read depth that could not reliably distinguish CN1 vs. CN2 when only one haplotype was found.

We next applied Paraphase to our collection of population samples (See Methods). While the sample sizes for non-European populations are small, among 259 unrelated European individuals, there are 6 (2.32%, all validated with MLPA) with one copy of *SMN1* (SMA 1+0 carriers), and 61 (23.6%) samples have *SMN2Δ7–8*, agreeing with previous studies^5,6,21^.

### *SMN1* and *SMN2* haplotypes across populations

We performed a population-wide haplotype analysis of 925 *SMN1* haplotypes and 645 *SMN2* haplotypes (excluding *SMN2Δ7–8*), and identified ten and nine major *SMN1* and *SMN2* haplogroups, respectively (Figure 2). Representative haplotype sequences from each haplogroup are shown in Figure 3, together with *SMN2Δ7–8* sequences. Through pedigree-based analysis (see Methods), we phased *SMN1* haplotypes into alleles and summarized their population frequencies (Table 2, *SMN2* allele frequencies are listed in Table S4). A few *SMN1* haplotypes are named with suffix “c” to indicate that the downstream region of *SMN1* is similar to that of *SMN2* (Figure 3). For example, S1-1c is similar to its corresponding haplotype without the suffix, S1-1, in the gene body and is similar to *SMN2* downstream of the gene. These haplotypes form separate clades and group with *SMN2* haplotypes when sequences of the upstream and downstream regions are included in the phylogenetic analysis (Figure S1A). These haplotypes could have arisen through gene conversion^33,34^.

**Table 2.** *SMN1* allele frequencies across five ethnic populations.

| *SMN1* Alleles | European |  | East Asian |  | South Asian |  | Admix American |  | African |  |
| --- | --- | --- | --- | --- | --- | --- | --- | --- | --- | --- |
| **Zero-copy (no *SMN1*)** | **5** | **1.2%** | **1** | **2.4%** | **1** | **1.9%** | **0** | **0.0%** | **0** | **0.0%** |
| **Singleton *SMN1* Alleles** |  |  |  |  |  |  |  |  |  |  |
| S1-1 | 233 | 55.9% | 35 | 83.3% | 27 | 51.9% | 47 | 67.1% | 26 | 29.9% |
| S1-1c | 16 | 3.8% | 1 | 2.4% | 2 | 3.8% | 2 | 2.9% | 2 | 2.3% |
| S1-2 | 80 | 19.2% | 2 | 4.8% | 7 | 13.5% | 6 | 8.6% | 1 | 1.1% |
| S1-2c | 0 | 0.0% | 1 | 2.4% | 1 | 1.9% | 0 | 0.0% | 0 | 0.0% |
| S1-3 | 65 | 15.6% | 0 | 0.0% | 7 | 13.5% | 8 | 11.4% | 1 | 1.1% |
| S1-4c | 7 | 1.7% | 0 | 0.0% | 1 | 1.9% | 1 | 1.4% | 0 | 0.0% |
| S1-5 | 1 | 0.2% | 0 | 0.0% | 0 | 0.0% | 1 | 1.4% | 1 | 1.1% |
| S1-6 | 0 | 0.0% | 0 | 0.0% | 0 | 0.0% | 1 | 1.4% | 3 | 3.4% |
| S1-6c | 0 | 0.0% | 0 | 0.0% | 0 | 0.0% | 1 | 1.4% | 1 | 1.1% |
| S1-7 | 0 | 0.0% | 0 | 0.0% | 0 | 0.0% | 0 | 0.0% | 2 | 2.3% |
| S1-8 | 0 | 0.0% | 0 | 0.0% | 0 | 0.0% | 0 | 0.0% | 1 | 1.1% |
| S1-9d | 0 | 0.0% | 0 | 0.0% | 0 | 0.0% | 0 | 0.0% | 1 | 1.1% |
| S1-9 | 0 | 0.0% | 0 | 0.0% | 0 | 0.0% | 0 | 0.0% | 9 | 10.3% |
| S1-10 | 0 | 0.0% | 0 | 0.0% | 0 | 0.0% | 0 | 0.0% | 8 | 9.2% |
| **Singleton total** | **402** | **96.4%** | **39** | **92.9%** | **45** | **86.5%** | **67** | **95.7%** | **56** | **64.4%** |
| **Two-copy *SMN1* Alleles** |  |  |  |  |  |  |  |  |  |  |
| S1-1+S1-1 | 3 | 0.7% | 0 | 0.0% | 2 | 3.8% | 0 | 0.0% | 0 | 0.0% |
| S1-1+S1-2 | 1 | 0.2% | 1 | 2.4% | 1 | 1.9% | 0 | 0.0% | 0 | 0.0% |
| S1-1+S1-3 | 4 | 1.0% | 0 | 0.0% | 0 | 0.0% | 0 | 0.0% | 0 | 0.0% |
| S1-1+S1-1c | 0 | 0.0% | 1 | 2.4% | 3 | 5.8% | 1 | 1.4% | 0 | 0.0% |
| S1-2+S1-2c | 1 | 0.2% | 0 | 0.0% | 0 | 0.0% | 0 | 0.0% | 0 | 0.0% |
| S1-4c+S1-4c | 1 | 0.2% | 0 | 0.0% | 0 | 0.0% | 0 | 0.0% | 0 | 0.0% |
| S1-5+S1-5 | 0 | 0.0% | 0 | 0.0% | 0 | 0.0% | 1 | 1.4% | 0 | 0.0% |
| S1-6+S1-6c | 0 | 0.0% | 0 | 0.0% | 0 | 0.0% | 1 | 1.4% | 1 | 1.1% |
| S1-8+S1-9d | 0 | 0.0% | 0 | 0.0% | 0 | 0.0% | 0 | 0.0% | 21 | 24.1% |
| S1-8+S1-9c | 0 | 0.0% | 0 | 0.0% | 0 | 0.0% | 0 | 0.0% | 2 | 2.3% |
| S1-9+S1-9 | 0 | 0.0% | 0 | 0.0% | 0 | 0.0% | 0 | 0.0% | 2 | 2.3% |
| S1-10+S1-10 | 0 | 0.0% | 0 | 0.0% | 0 | 0.0% | 0 | 0.0% | 2 | 2.3% |
| S1-1+S1-9d | 0 | 0.0% | 0 | 0.0% | 0 | 0.0% | 0 | 0.0% | 2 | 2.3% |
| S1-1+S1-8 | 0 | 0.0% | 0 | 0.0% | 0 | 0.0% | 0 | 0.0% | 1 | 1.1% |
| **Two-copy total** | **10** | **2.4%** | **2** | **4.8%** | **6** | **11.5%** | **3** | **4.3%** | **31** | **35.6%** |
| Total alleles | 417 |  | 42 |  | 52 |  | 70 |  | 87 |  |

For single-copy *SMN1* alleles, S1-1 is the most common haplotype across all ethnicities, with a frequency ranging from 29.9% in Africans to 83.3% in East Asians. S1-2 and S1-3 are also common (10-20%) in Europeans, South Asians and Admixed Americans, while they are less common (<3%) in Africans and East-Asians. Notably, it is not the most common haplotype, S1-1, but S1-2 that is represented by the reference genome (GRCh38). Additionally, we observed several African-specific haplogroups (S1-7, S1-8, S1-9, S1-9d and S1-10). Out of all *SMN1* haplogroups, S1-10 is closest in sequence to *SMN2* (Figure 2, Figure 3 and Figure S2).

The sequence differences between *SMN1* and *SMN2* are mainly located in Exon 7 and Exon 8, as well as the downstream region (Figure 3). *SMN2Δ7–8* is a truncated form with a 6.3kb deletion of Exons 7-8^21,23^ (Figure 3), and its downstream region is highly similar to that of *SMN2*, confirming previous findings that this common truncated form likely derives from *SMN2*^21^. Conversely, the upstream region and Exons 1-6 are highly similar between *SMN1* and *SMN2* and there is not a single SNP that could distinguish *SMN1* from *SMN2* reliably in this region, i.e. there is not any SNP that is present in <10% of *SMN1* haplotypes and >90% of *SMN2* haplotypes, or vice versa. *SMN1* and *SMN2* haplotypes do not separate when only Exons 1-6 sequences are included in the phylogenetic analysis (Figure S1B). As a result of the high similarity, read alignments are often ambiguous in this region, even for long reads.

In addition to small variants and the 6.3kb known deletion in *SMN2*, we also found a previously unknown common structural variant in this region. A 3.6kb (chr5:70917700-70921260, GRCh38) deletion occurs upstream of *SMN1* in S1-9d, which is otherwise similar to S1-9.

### Two-copy *SMN1* alleles

African individuals have more copies of *SMN1* than other populations, with about 45-50% of the population carrying >2 copies of *SMN1* indicating the presence of two-copy *SMN1* alleles^5,6,21^. The higher frequency of two-copy *SMN1* alleles leads to a higher frequency (estimated at ∼27% of all carriers) of 2+0 silent carriers where an individual has two copies of *SMN1* but both occur on the same chromosome. Without pedigree information, two-copy *SMN1* alleles are impossible to detect directly with current technologies. Through pedigree-based phasing of haplotypes into alleles, we studied two-copy *SMN1* alleles and their frequencies. In African individuals, there exist a few haplotypes (S1-8, S1-9c and S1-9d) that are commonly found in two-copy *SMN1* alleles but not singleton *SMN1* alleles (Table 2) and these could serve as potential markers for two-copy *SMN1* alleles. In particular, we identified a common two-copy *SMN1* allele, S1-8+S1-9d, that comprises two thirds (21 out of 31) of African two-copy *SMN1* alleles and 24.1% of total African alleles. These two *SMN1* haplotypes, S1-8 and S1-9d, are rarely present as singletons (both at 1.1%, Table 2). Taking the previous estimate of zero-copy *SMN1* allele frequency in Africans (0.68%^6^), if an African individual has two copies of *SMN1*, S1-8 and S1-9d, the likelihood of the two haplotypes being on the same chromosome, i.e. a silent carrier (2+0), is 7.7 times higher than the two haplotypes being present on different chromosomes, and thus the probability of being a silent carrier is 88.5%.

The SNP g.27134T>G in Intron 7 of *SMN1* is commonly used as a marker of two-copy *SMN1* alleles^24^. In our data, this SNP is only found in haplogroups S1-8 (21.9%), S1-9 (100%), S1-9c (100%) and S1-9d (96.3%). Samples positive for g.27134T>G are mainly those carrying the two-copy alleles S1-8+S1-9d, S1-8+S1-9c and S1-9 singletons. S1-9 is commonly found as singleton *SMN1* alleles in Africans (10.3% of all African alleles and 16.1% of singleton African alleles) and it differs from S1-9d only by the 3.6kb deletion upstream of *SMN1* and differs from S1-9c only in the downstream region. Therefore, g.27134T>G is expected to be present on a high percentage of singleton *SMN1* alleles (16.1% in our data), consistent with previous maximum-likelihood estimates (18.4%)^21^, and thus not an accurate marker for two-copy *SMN1* alleles. Conversely, using HiFi reads, Paraphase can accurately distinguish S1-9d or S1-9c from S1-9. In addition, being able to identify the other haplotype of the pair, S1-8, further improves Paraphase’s accuracy of detecting the two-copy *SMN1* alleles.

For non-African populations, 57.1% (12 of 21) of two-copy *SMN1* alleles involve combinations of common *SMN1* haplotypes, i.e. S1-1+S1-1, S1-1+S1-2 and S1-1+S1-3 (Table 2). We also observed four two-copy *SMN1* alleles where one of the copies of *SMN1* includes the *SMN2* sequence in the downstream region (flagged with the “c” suffix), i.e. S1-1+S1-1c, S2-2+S2-2c, S1-4c+S1-4c and S1-6+S1-6c. It is possible that gene conversion from *SMN2* to *SMN1* in Exons 7-8 resulted in these two-copy *SMN1* alleles. Taking all non-African samples together, this pattern explains 8 (38.1%) out of 21 two-copy *SMN1* alleles, or 4 (50%) out of 8 distinct two-copy *SMN1* alleles. This is in line with the previous finding that paralog specific variants (PSVs) between *SMN1* and *SMN2* downstream of the genes are overrepresented in signature variants enriched in two-copy *SMN1* alleles in a Chinese population^25^. However, as these “c” haplotypes are also present as singleton *SMN1* alleles (6.0% of all non-African singleton alleles) and the other haplotype of the pair is often a highly common singleton allele such as S1-1 and S1-2, these haplotypes will frequently occur on two different chromosomes, so this “c” haplotype pattern as a marker does not have a high PPV as was observed for the S1-8+S1-9d allele in Africans.

### Co-segregation of *SMN1* and *SMN2* haplotypes

We next investigated the co-segregation of *SMN1* and *SMN2* haplotypes. Our results show that *SMN2* (including *SMN2Δ7–8*) is present on 85.3% of singleton *SMN1* alleles but only 26.9% of two-copy *SMN1* alleles. This indicates that gains of *SMN1* are often accompanied with losses of *SMN2*^21^ and it is possible that many two-copy *SMN1* alleles were generated through gene conversion of *SMN2* into *SMN1*^33^.

For standard alleles with one copy of *SMN1* and one copy of full-length *SMN2*, i.e. excluding *SMN2Δ7– 8*, we examined the types of *SMN1* and *SMN2* haplotypes on the same allele. We found that an *SMN1* haplogroup is usually segregated with a specific *SMN2* haplogroup (Table 3). This suggests that it is possible to probabilistically phase *SMN1* and *SMN2* together. For simplicity we named the *SMN2* haplogroups to match the corresponding *SMN1* haplogroups that they usually co-segregate with (e.g. S1-1 and S2-1 usually co-segregate). Interestingly, when we queried the sequence similarity between *SMN1* and *SMN2* haplogroups in Exons 1-6 (Exons 7-8 are not included as they are differentiated between *SMN1* and *SMN2*), *SMN1* haplogroups usually share the highest similarity with the co-segregating *SMN2* haplogroups (Figure S3A). This is true for the three most common haplogroups (S1-1, S1-2 and S1-3), as well as 3 out of the 6 less common haplogroups (S1-4 through S1-9, S1-10 is not included as none of S1-10 haplotypes occurs on the same allele as *SMN2*, see below). As a result, some of the co-segregating *SMN1* and *SMN2* haplogroups group together when Exons 1-6 sequences were used to create the phylogeny (Figure S1B). For less common alleles, a larger sample size is needed to further confirm the co-segregation pattern and the sequence similarity, especially for S1-7 (N=2) and S1-8 (N=1).

**Table 3.** *SMN1*-*SMN2* haplogroup co-segregation on alleles with one copy of full-length *SMN1* and one copy of full-length *SMN2*.

| *SMN1* haplogroup | *SMN2* haplogroup | # co-segregated alleles | # *SMN1* haplogroups segregated with other *SMN2* haplogroups | # *SMN2* haplogroups segregated with other *SMN1* haplogroups | % co-segregation |
| --- | --- | --- | --- | --- | --- |
| S1-1/S1-1c | S2-1 | 297 | 8<sup>a</sup> | 8<sup>b</sup> | 94.9% |
| S1-2/S1-2c | S2-2 | 101 | 0 | 2 | 98.1% |
| S1-3 | S2-3 | 70 | 5<sup>b</sup> | 6<sup>a</sup> | 86.4% |
| S1-4c | S2-4 | 8 | 2 | 0 | 80.0% |
| S1-5 | S2-5 | 3 | 0 | 0 | 100.0% |
| S1-6/S1-6c | S2-6 | 4 | 1 | 0 | 80.0% |
| S1-7 | S2-7 | 2 | 0 | 0 | 100.0% |
| S1-8 | S2-8 | 1 | 0 | 0 | 100.0% |
| S1-9/S1-9d | S2-9 | 8 | 0 | 0 | 100.0% |

a. Among these alleles, 6 are S1-1 co-segregated with S2-3.
b. Among these alleles, 5 are S1-3 co-segregated with S2-1.

We also examined co-segregation of alleles other than one copy of *SMN1* and one copy of full-length *SMN2*. First, S1-10 alleles always contain zero copy of *SMN2* (8 out of 8 alleles). Since S1-10 is closest in sequence to *SMN2* among all *SMN1* haplogroups (Figure 2) and S1-10 alleles never contain *SMN2*, S1-10 could be a hybrid gene between *SMN1* and *SMN2* created by a fusion deletion. Next, *SMN2Δ7–8* alleles segregate with S1-1 in 98% (51 out of 52) of cases. *SMN2Δ7–8* is most similar in sequence in Exons 1-6 to S1-1 and S2-1 (Figure S3B). Both the co-segregation and the sequence similarity suggest that *SMN2Δ7–8* is most likely derived from S2-1. Finally, we summarized the frequency of *SMN1* (*SMN2*) haplotypes on alleles without *SMN2* (*SMN1*) (Table S5). Among our limited sample of 9 alleles without *SMN1* (zero-copy *SMN1* alleles), four contain more than one copy of *SMN2*. Among these four alleles, two of them carry an *SMN2* haplotype with the downstream region similar to *SMN1* (Figure S4), suggesting possible loss of *SMN1* through gene conversion from *SMN1* to *SMN2*.

## Discussion

Here we provide the most comprehensive analysis of variation in one of the most difficult, clinically important regions of the human genome. Extending beyond copy number testing based primarily on c.840C>T as is often done, Paraphase phases the region to provide a much richer level of information. Using the phasing information, Paraphase can detect other pathogenic variants and enable haplotype-based screening of silent carriers. Since Paraphase works mainly by assembling variant positions from long reads, it works for both WGS and hybrid capture-based enrichment data, and can be adapted to work with amplicon sequencing data, when the full *SMN1*/*SMN2* regions are captured or amplified. Compared with short-read based methods, highly accurate HiFi reads can provide long-range haplotype information through entire genes and easily pick up large structural variants such as the 6.3kb deletion in *SMN2Δ7–8* and the 3.6kb deletion in the *SMN1* haplotype S1-9d.

In this study we conducted, to our knowledge, the first population-wide full-gene haplotype analysis of *SMN1* and *SMN2*. Combining our gene level phasing with pedigree information, we identified haplotypes that form two-copy *SMN1* alleles. Most importantly, we identified a common two-copy *SMN1* allele that comprises 67.7% of two-copy *SMN1* alleles in Africans. The two individual haplotypes on this allele each occur very rarely as singleton *SMN1* alleles in the population. Based on our limited sample of 87 African alleles, we estimate that testing positive for these two haplotypes in an individual with two copies of *SMN1* gives a silent carrier risk of 88.5%, which is significantly higher than the previously found marker SNP g.27134T>G (1.7-3.0%)^20,21,24^.

In addition, we found co-segregation patterns between *SMN1* and *SMN2* haplotypes. An *SMN1* haplogroup often co-segregates with the *SMN2* haplogroup that is most similar in sequence, suggesting that intrachromosomal gene conversion between *SMN1* and *SMN2* plays a significant role in the evolution of this region. With larger sample datasets enabling more accurate allele frequency calculations, it should be possible to build a probabilistic model to predict the most likely allele/genotype configurations based on the haplotypes seen in an individual. This would be very helpful for silent carrier detection. For example, an individual with S1-8, S1-9d and S2-1 is very likely a silent carrier, as S1-8 and S1-9d rarely exist as singleton *SMN1* alleles and S2-1 rarely segregates with S1-8 or S1-9d. For an individual with these haplotypes, the most likely alleles are two copies of *SMN1* (S1-8+S1-9d) with no *SMN2* on one allele and one copy of *SMN2* (S2-1) with no *SMN1* on the other allele.

One limitation in this study is the relatively small number of samples (438) studied. To make more statistically powered findings out of the haplotype analysis, it is desirable to increase the sample size, particularly for non-European populations. Future analysis of large population data with Paraphase, using either HiFi WGS or possibly a hybrid capture based or other targeted long-read approaches, will allow a better characterization of variants in both genes, identification of more diverse haplotypes and more genetic markers for silent carrier detection.

The method employed in Paraphase can be applied to other segmental duplication regions with extremely high sequence similarity and frequent copy number variations. We are currently extending this method to solve similar gene paralog problems such as *CYP21A2*, and will apply this method to more clinically relevant genes in the future. The development of more targeted informatics solutions for difficult regions with HiFi data will bring us one step closer to consolidating the numerous genetic tests that are currently offered into a single test.

## Data and code availability

Paraphase can be downloaded from https://github.com/PacificBiosciences/paraphase.

Bamlets for visualizing *SMN1* and *SMN2* haplotypes of AGBT and HPRC samples can be downloaded from https://github.com/xiao-chen-xc/SMN_phased_data.

## Acknowledgements

We thank the Human Pangenome Reference Center (HPRC) for generating and releasing the HiFi WGS data. This research was made possible through access to the data and findings generated by the 100,000 Genomes Project. The 100,000 Genomes Project is managed by Genomics England Limited (a wholly owned company of the Department of Health and Social Care). The 100,000 Genomes Project is funded by the National Institute for Health Research and NHS England. The Wellcome Trust, Cancer Research UK and the Medical Research Council have also funded research infrastructure. The 100,000 Genomes Project uses data provided by patients and collected by the National Health Service as part of their care and support. The GA4K HiFi sequencing data was made possible by the generous gifts to Children’s Mercy Research Institute and the Genomic Answers for Kids program at Children’s Mercy Kansas City.

## Declaration of interest

X.C., J.H. and M.A.E. are employees of Pacific Biosciences.

## Notes

### Summary of Updates

Supplemental files updated.

