## Supplemental text, figures and tables for "Comprehensive *SMN1* and *SMN2* profiling for spinal muscular atrophy analysis using long-read PacBio HiFi sequencing"

### Supplemental Information

#### Variability of paralog specific variants (PSVs) between *SMN1* and *SMN2*

Previous studies^1,2^ using short-read population data analyzed the paralog specific variants (PSVs) between *SMN1* and *SMN2* of the reference genome and found indirectly (i.e. without phasing) that they are much more variable in African populations than non-African populations. This calls for careful selection of PSVs for short read-based *SMN1/SMN2* copy number calculation^1,2^. Here we analyzed the variability of these PSVs on our phased *SMN1* and *SMN2* haplotypes. While PSVs are mostly fixed in non-African populations, African haplotypes show a much higher rate of sharing - i.e. *SMN2* bases in an *SMN1* haplotype, or *SMN1* bases in an *SMN2* haplotype (Figure S2A). Focusing on 15 reference-PSVs flanking c.840 in Intron 6-Exon 8, 33.5% of African haplotypes have at least 2 discrepant sites (13.3% have at least 5 discrepant sites), while most (96%) non-African haplotypes have zero or one (Figure S2A). The biggest contributors to the high PSV discrepancy in Africans are a few African-specific haplogroups (Figure S2B and Figure S2C), such as S1-10 (7 or more discrepant sites), and S2-9 (5 discrepant sites).

#### Silent carrier risk calculation of S1-8+S1-9d in Africans

We took the frequency of S1-8+S1-9d (21/31) out of two-copy *SMN1* alleles, as well as the frequency of S1-8 (1/56) and S1-9d (1/56) out of singleton *SMN1* alleles from our data. We took the frequency of zero-copy (0.68%), singleton (71.79%) and two-copy (27.51%) *SMN1* alleles from Sugarman et al^3^. The probability of S1-8/S1-9d is 2*(71.79%*1/56)*(71.79%*1/56). The probability of -/S1-8+S1-9d is 2*0.68%*(27.51%*21/31). The silent carrier risk is calculated as the weighted probability of -/S1-8+S1-9d.

#### *SMN1*/*SMN2* variant calls

The *SMN1*/*SMN2* gene is 27.9kb long and consists of 8 exons, among which Exon 1 is far away from the rest of the exons (13.7kb away from Exon 2). Due to the distance and the fact that *SMN1* and *SMN2* are highly similar in sequence in Exons 1-6, it could be more challenging to phase haplotypes through Exon 1 than Exons 2-8. Among the haplotypes resolved by Paraphase, 98.5% of them cover Exons 2-8, and 88.4% of them cover Exons 1-8. Note that Exon 1 encodes 27 amino acids and currently there is only one pathogenic/likely pathogenic variant in Exon 1 with more than one star in ClinVar (ClinVar ID:9168) (ClinVar last accessed on Oct 12, 2022).

Small variants were called in each assembled haplotype using DeepVariant. Among the protein changing variants in *SMN1*, we identified two missense variants and one in-frame insertion. They are:

S4G, 70925113A>G, not in ClinVar

G6S, 70925119G>A, not in ClinVar

G7GSGGGV, 70925123G>GCAGTGGTGGCGGCGT, not in ClinVar

K93T, 70942362A>C, ClinVar ID:638580, uncertain significance

Among the protein changing variants in *SMN2*, we identified three missense variants. They are:

G26D, 70049762G>A, not in ClinVar

G106S, 70066976G>A, not in ClinVar

G287R, 70076545G>C, called in four samples. This variant was previously shown to be a positive modifier of SMA^4^.

Interestingly, G106S is reported for *SMN1* in ClinVar (ID:634938, uncertain significance), and G26D has been reported by a previous study^5^ where they identified the variant but could not map it to *SMN1* or *SMN2*. It is possible that these variants can occur on either *SMN1* or *SMN2*, or these are *SMN2*-specific variants that were mapped to *SMN1* by mistake in the case of G106S.

#### Supplementary figures

##### Figure S1. Trees of the same set of haplotypes used in Figure 2 created with gene sequences plus upstream/downstream regions (A) and Exons 1-6 only (B).

Haplogroups are colored in the same way as in Figure 2. In Panel B, shaded nodes indicate *SMN2* haplogroups. Some *SMN1* and *SMN2* haplogroups of the same color (co-segregating haplogroups) group together (green, purple, blue, magenta and orange, etc.). The inset shows the same tree reduced to two colors (red: *SMN1*; black: *SMN2*).

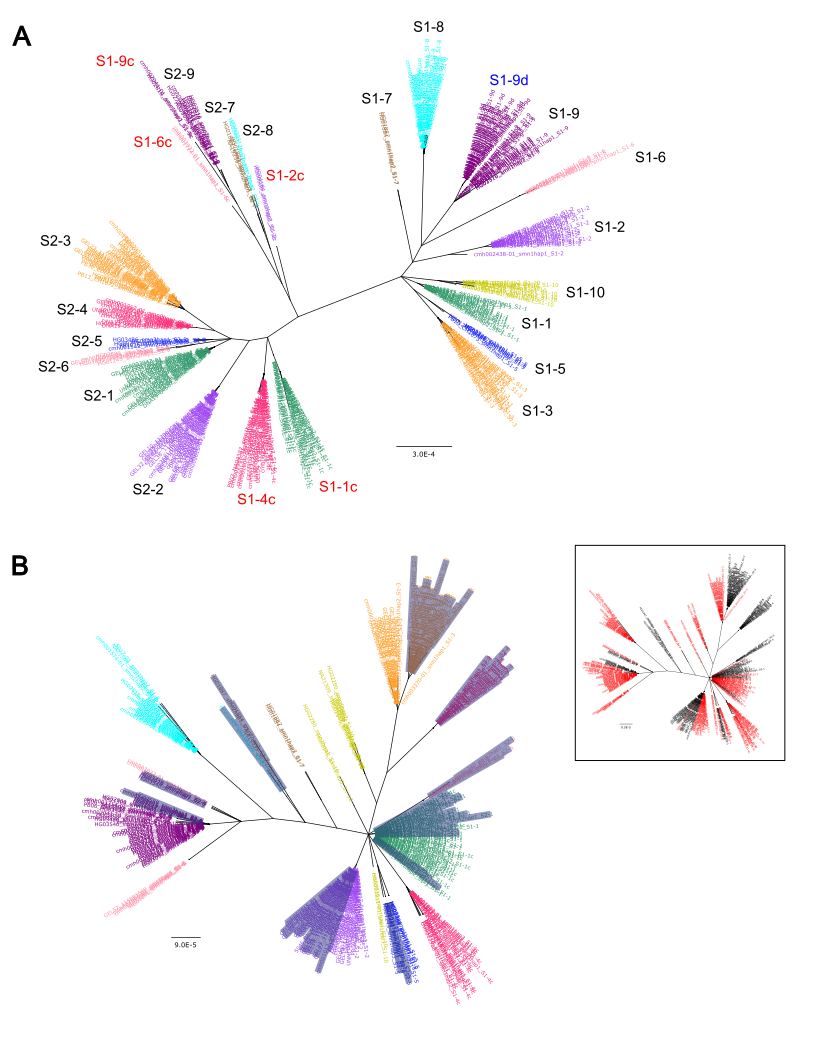

##### Figure S2. Discrepant PSV sites across populations.

**A.** Frequency of haplotypes carrying discrepant sites across populations. The x axis shows the number of discrepant PSV sites, i.e. *SMN2* bases on *SMN1* haplotypes or *SMN1* bases on *SMN2* haplotypes, out of 15 reference-PSVs flanking c.840C, taken from Chen et al. 2020^1^. **B.** Frequency of haplotypes carrying discrepant sites across *SMN1* haplotypes. The “c” and “d” haplotypes are identical to their corresponding haplotypes in the gene body, so they are considered as their corresponding haplotypes, e.g. S1-1c considered as S1-1. **C.** Frequency of haplotypes carrying discrepant sites across *SMN2* haplotypes.

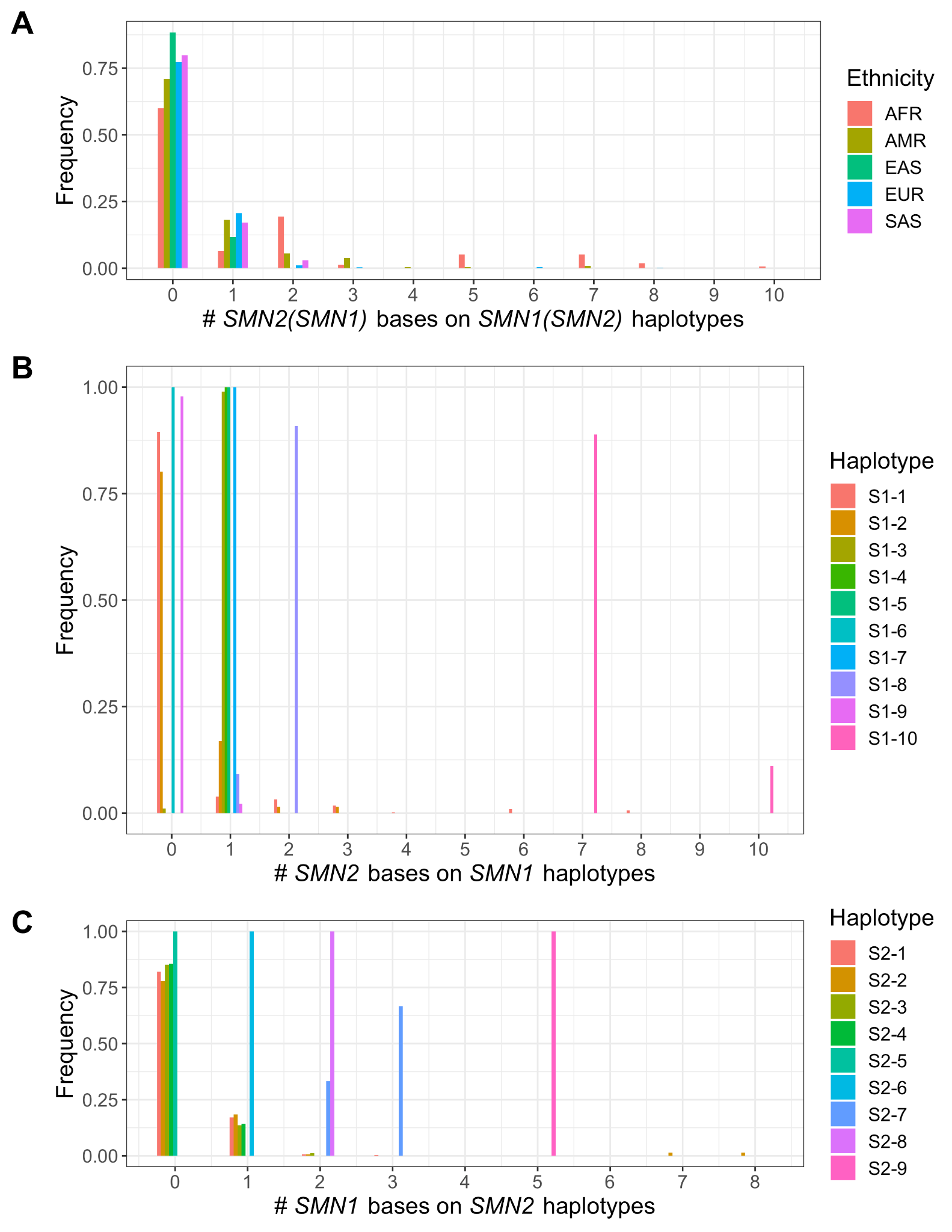

##### Figure S3. Sequence similarity between *SMN1* and *SMN2* haplogroups.

**A.** *SMN1* haplotypes are compared against *SMN2* haplotypes and the weighted average similarity between each haplogroup is plotted. For each pairwise comparison, variant concordance is calculated as the fraction of concordant bases out of 444 total sites where variants occur across populations in Exons 1-6. The “c” and “d” haplotypes are identical to their corresponding haplotypes in Exons 1-6, so they are considered as their corresponding haplotypes, e.g. S1-1c considered as S1-1. **B.** *SMN2∆7–8* haplotypes are compared against *SMN1* and *SMN2* haplotypes among the same set of 444 total variant sites in Exons 1-6. Variant concordance calculation is the same as in A.

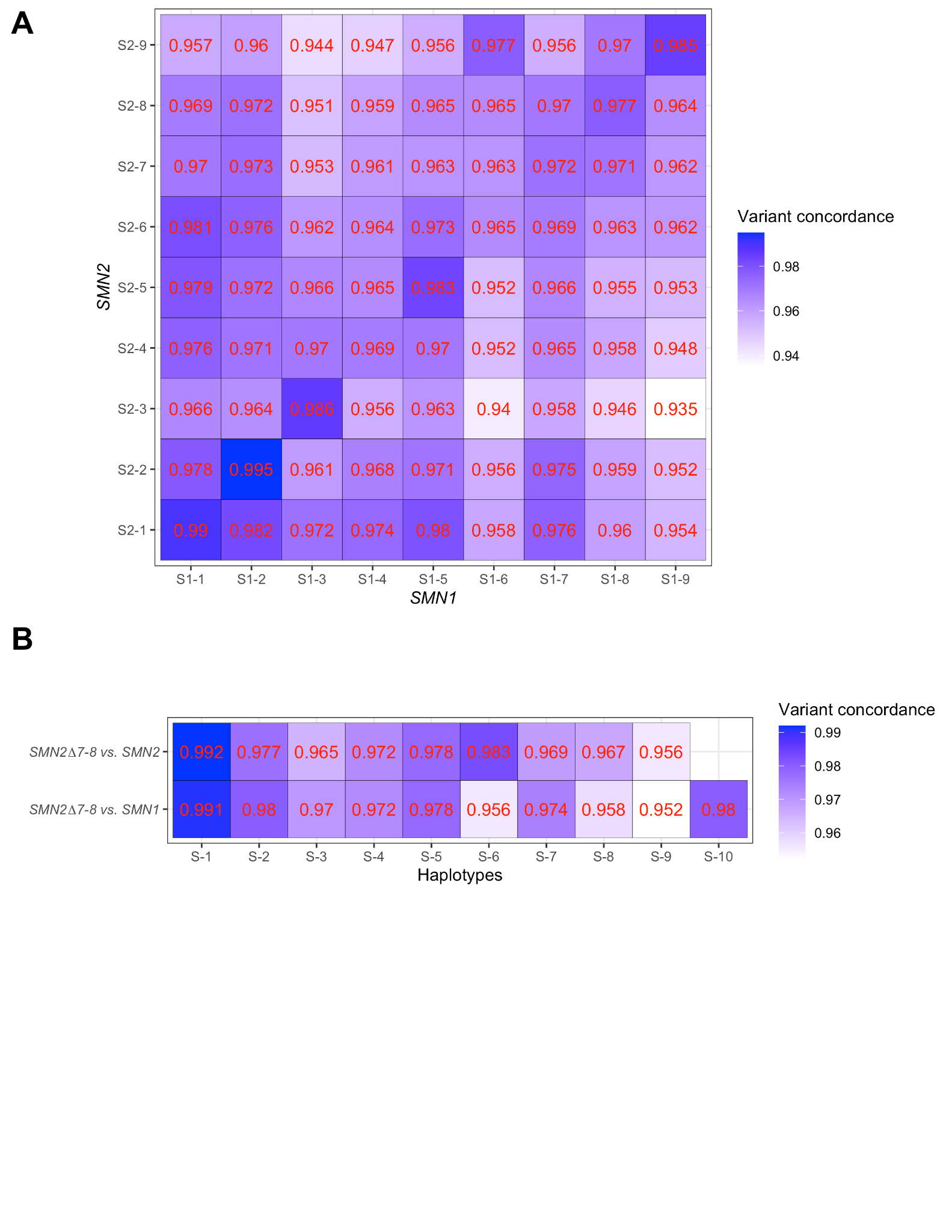

##### Figure S4. IGV snapshot of *SMN2* haplotypes with the downstream region similar to *SMN1.*

In HG02132, the downstream region of *SMN2* haplotype 3 is similar to *SMN1*. In GEL02, the downstream region of *SMN2* haplotype 4 is similar to *SMN1*. Reads in blue are uniquely assigned to a haplotype, while reads in gray can be assigned to more than one possible haplotype and a random one is selected (this happens when haplotypes are identical over a region).

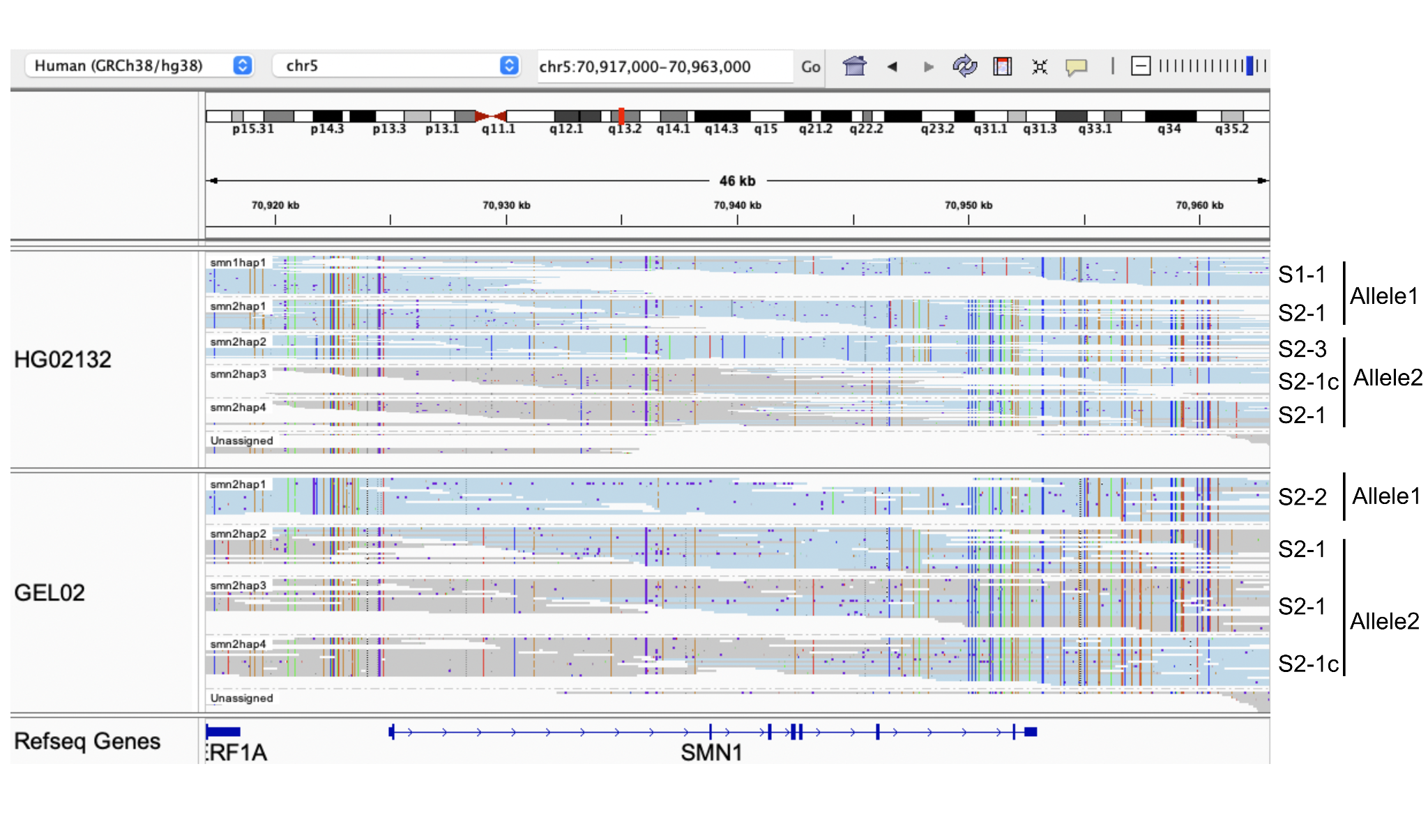

#### Supplementary tables

##### Table S1. Validation sample details. (Excel Spreadsheet)

##### Table S2. Pedigree information.

| Data source | EUR | AFR | EAS | SAS | AMR | unknown | mixed ancestry | notes |
| --- | --- | --- | --- | --- | --- | --- | --- | --- |
| RadboudUMC  (Kucuk et al. in review^6^) | 8 | 0 | 0 | 0 | 0 | 0 | 0 | 30X HiFi WGS for all samples |
| 100,000 Genomes Project (GEL) | 1 | 0 | 0 | 0 | 0 | 0 | 0 | 30X HiFi WGS for all samples |
| GIAB | 1 | 0 | 1 | 0 | 0 | 0 | 0 | 30X HiFi WGS for all samples |
| ChineseQuartet | 0 | 0 | 1 | 0 | 0 | 0 | 0 | 30X HiFi WGS for all samples |
| HPRC/1kGP | 0 | 29 | 16 | 24 | 28 | 0 | 0 | 30X HiFi WGS genomes for the proband and  30X short read WGS data for the parents |
| GA4K | 188 | 8 | 0 | 2 | 7 | 9 | 18 | 20-30X HiFi WGS genomes for the proband  and 5-10X HiFi genomes for the parents |
| Total | 198 | 37 | 18 | 26 | 35 | 9 | 18 |  |

##### Table S3. Population sample results. (Excel Spreadsheet)

##### Table S4. *SMN2* allele frequencies across five ethnic populations.

|  | EUR | | EAS | | SAS | | AMR | | AFR | |
| --- | --- | --- | --- | --- | --- | --- | --- | --- | --- | --- |
| no *SMN2* | 54 | 12.9% | 5 | 11.9% | 13 | 25.0% | 12 | 17.1% | 43 | 49.4% |
| S2-1 | 163 | 39.1% | 33 | 78.6% | 25 | 48.1% | 38 | 54.3% | 27 | 31.0% |
| S2-2 | 80 | 19.2% | 3 | 7.1% | 7 | 13.5% | 5 | 7.1% | 1 | 1.1% |
| S2-3 | 61 | 14.6% | 0 | 0.0% | 7 | 13.5% | 4 | 5.7% | 1 | 1.1% |
| S2-4 | 7 | 1.7% | 0 | 0.0% | 0 | 0.0% | 1 | 1.4% | 0 | 0.0% |
| S2-5 | 1 | 0.2% | 0 | 0.0% | 0 | 0.0% | 2 | 2.9% | 1 | 1.1% |
| S2-6 | 0 | 0.0% | 0 | 0.0% | 0 | 0.0% | 2 | 2.9% | 2 | 2.3% |
| S2-7 | 0 | 0.0% | 0 | 0.0% | 0 | 0.0% | 0 | 0.0% | 2 | 2.3% |
| S2-8 | 0 | 0.0% | 0 | 0.0% | 0 | 0.0% | 0 | 0.0% | 1 | 1.1% |
| S2-9 | 0 | 0.0% | 0 | 0.0% | 0 | 0.0% | 0 | 0.0% | 8 | 9.2% |
| *SMN2∆7–8* | 44 | 10.6% | 0 | 0.0% | 0 | 0.0% | 5 | 7.1% | 0 | 0.0% |
| more than one copy of *SMN2* | 7 | 1.7% | 1 | 2.4% | 0 | 0.0% | 1 | 1.4% | 1 | 1.1% |
| Total | 417 |  | 42 |  | 52 |  | 70 |  | 87 |  |

##### Table S5. Pan-ethnic frequencies of *SMN1 (SMN2)* haplotypes on alleles without *SMN2 (SMN1)*.

| *SMN1* | *SMN2* | # alleles | percentage |
| --- | --- | --- | --- |
| S1-1 | no *SMN2* | 64 | 47.1% |
| S1-2 |  | 11 | 8.1% |
| S1-3 |  | 11 | 8.1% |
| S1-6 |  | 1 | 0.7% |
| S1-9 |  | 3 | 2.2% |
| S1-10 |  | 8 | 5.9% |
| two copies of *SMN1* |  | 38 | 27.9% |
| Total |  | 136 |  |
| no *SMN1* | S2-1 | 1 | 11.1% |
|  | S2-2 | 4 | 44.4% |
|  | S2-2+S2-2 | 1 | 11.1% |
|  | *SMN2*∆7–8+S2-2 | 1 | 11.1% |
|  | S2-1+S2-1+S2-1c | 1 | 11.1% |
|  | S2-3+S2-1+S2-1c | 1 | 11.1% |
|  | Total | 9 |  |
